# NFAT signaling is indispensable for persistent memory responses of MCMV-specific CD8^+^ T cells

**DOI:** 10.1101/2023.05.02.539029

**Authors:** M. Zeeshan Chaudhry, Lisa Borkner, Friederike Berberich-Siebelt, Luka Cicin-Sain

## Abstract

Cytomegalovirus (CMV) induces a unique T-cell response, where antigen-specific populations do not contract, but rather inflate during viral latency. It has been proposed that subclinical episodes of virus reactivation feed the inflation of CMV-specific memory cells by intermittently engaging T-cell receptors (TCRs), but evidence of TCR engagement has remained lacking. Nuclear factor of activated T cells (NFAT) is a family of transcription factors, where NFATc1 and NFATc2 signal downstream of TCR in mature T lymphocytes. We show selective impacts of NFATc1 and/or NFATc2 genetic ablations on the long-term inflation of MCMV-specific CD8 T-cell responses despite largely maintained responses to acute infection. NFATc1 ablation elicited robust phenotypes in isolation, but the strongest effects were observed when both NFAT genes were missing. CMV control was impaired only when both NFATs were deleted in CD8 T cells used in adoptive immunotherapy of immunodeficient mice. Transcriptome analyses revealed that T-cell intrinsic NFAT is not necessary for CD8 T-cell priming, but rather for their maturation towards effector-memory and in particular the effector cells, which dominate the pool of inflationary cells.

## INTRODUCTION

The development of immune memory to viruses is usually studied in the context of primary virus infections, which lead to a potent immune response that clears virus infection. This immune memory protects the host during reinfection and generates an anamnestic response upon secondary infections (1, 2). Chronic viruses, on the other hand, have developed immune evasion strategies that allow them to persistently maintain themselves in the host (3-5). Viral persistence can induce chronic inflammatory conditions and exhaust the adaptive host immune system over time, in particular the CD8+ T cell compartment (6). The exhaustion of virus-specific CD8+ T cells is characterized by poor effector cytotoxic activity, impaired cytokine production, and sustained expression of inhibitory receptors, such as programmed cell death-1 (PD-1)(7). In other cases, such as herpesvirus infection, the chronic virus infection is associated with long-term persistence of functional T cells despite continuous antigen stimulation (8-10).

Human cytomegalovirus (HCMV) is a prototypic herpesvirus that induces long-term chronic latent infection in human host. HCMV is of particular concern as a major cause of morbidity and mortality in transplant patients (11, 12). Normally in humans, the progressive expansion and persistence of virus-specific CD8^+^ T cells in the context of chronic CMV infection leads to a massive enrichment of an oligoclonal pool of CD45RA+CCR7- effector memory (TEMRA) cells. These cells lose the expression of classic memory markers like CD28 and IL-7R. This phenomenon is collectively referred to as ‘memory inflation’ (13). The murine CMV (MCMV) infection faithfully replicates the functional phenotype of inflationary T cell responses in mouse models and has been widely used to model human CMV infection (14).

CMV-specific persistent CD8^+^ T cells responses are maintained by continuous production of short-lived effector cells (SLECs) that are characterized by downregulation of costimulatory receptors CD28, CD27, and also of the IL-7 receptor alpha chain (CD127) (15, 16). These SLECs express high levels of KLRG1 and the NK cell-inhibitory molecule NKG2A (15). Another molecule expressed on inflationary cells is the chemokine receptor CX3CR1, a fractalkine receptor that is not expressed on conventional memory T cells generated following acute infections (17). Interestingly, conventional and inflationary CD8^+^ T cell responses emerge in parallel against different viral epitopes of CMV within the same host (16). This dichotomy is explained by differences in gene expression (18) and by the differential processing of persistent antigens, which require processing by the constitutive proteasome (19) in non-hematopoietic cells (20). The persistent CD8^+^ T cell responses are not induced by cross-presentation, but only if the target epitope is presented directly on the virus-infected cells (19). The conventional and inflationary CD8^+^ T cells also differ in their costimulatory requirements, where CD27 and CD28 costimulation is crucial for conventional CD8^+^ T cell responses but dispensable for persistent responses (21, 22). The lack of CD27 and CD28 costimulatory molecules on persistent CD8^+^ T cells and their stimulation by direct antigen presentation suggest that direct TCR ligation is decisive for these responses, but this has not been demonstrated.

T cell receptor (TCR) ligation by antigen and the resulting TCR-calcium-calcineurin axis represent a central signaling pathway for T cell activation. The TCR-mediated increase in intracellular Ca^2+^ and subsequent activation of calcineurin is responsible for *nuclear factor of activated T cells* (NFAT) signaling, i.e. its nuclear translocation and transactivation of target genes (23). NFAT family consists of five members first known as transcriptional regulators of T cell development, maturation and activation. Among these, NFATc1 to NFATc4 are activated by calcium-calcineurin signaling (24), of which only NFATc1 to NFATc3 are expressed in thymocytes and T cells. Among those, NFATc1 (also called NFAT2) and NFATc2 (NFAT1) play dominant roles in T cell activation and function. All NFAT members share a DNA binding domain that allows them to bind to a consensus DNA motif (25). In CD4^+^ T and Treg cells, different NFAT members have overlapping as well as distinct roles (26-29). In addition, NFATc1 and NFATc2 have been shown to regulate CD8^+^ T-cell differentiation following acute LCMV infection (30), whereas especially NFATc2 is important for chronic LCMV-specific CD8^+^ T cells responses and inducing CD8^+^ cell exhaustion (31). However, also the short isoform of NFATc1, NFATc1/αA (32) is highly upregulated during chronic LCMV infection (33). Despite this, the role of NFAT signaling or even a possible distinct role of NFATc1 versus NFATc2 in maintenance of persistent T-cells responses during chronic CMV infections has not been explored so far.

In this study, we characterized the role of NFAT signaling in CMV-specific responses during acute and chronic infection. We show that NFAT-deficient CD8^+^ T cells can mount robust CMV-specific responses following acute infection, but NFATc1 is indispensable for maintaining persistent CD8^+^ cell responses during the chronic phase. Furthermore, NFATc1 and NFATc2 differentially regulate the CD8^+^ cell transcriptome that leads to differences in T cell differentiation during acute and chronic infection phases. The loss of NFATc1 in virus-specific CD8^+^ T cells induces a memory phenotype and reduction in SLEC production. Moreover, the combined loss of NFATc1 and NFATc2 enforces a stronger memory phenotype in form of an accumulation of central memory T cells. Interestingly, despite defects in T cell activation and differentiation, CD8^+^ T cells with selective ablation of NFATc1 or NFATc2 efficiently controlled the virus in immunocompromised animals. However, CD8^+^ T cells with combined loss of NFATc1 and NFATc2 showed reduced antiviral capacity. These results strongly indicate that TCR-mediated NFAT signaling is crucial in maintaining persistent T cell accumulation during chronic virus infection.

## MATERIALS AND METHODS

### Mice

*Nfatc1^fl/fl^* (formerly called *Nfat2*^fl/fl^).*Cd4cre* (NFATc1 KO)*, Nfat2^−/−^* (NFATc2 KO), *Nfatc1^fl/fl^.Cd4cre. Nfat2^−/−^* (NFATc1c2 DKO) animals have been described before (31, 34, 35). C57BL/6-Thy1^a^- Tg(TcraTcrb)1100Mjb/Crl (CD90.1 OTI), B6.SJL-Ptprc^a^Pepc^b^/BoyJ (B6 CD45.1), C;129S4-Rag2^tm1.1Flv^- Il2rg^tm1.1Flv^/J (RAG2gc KO) and breeding pairs were originally purchased from Jackson Laboratory and bred under specific pathogen free conditions. NFAT KO animals were crossed with CD90.1 OTI mice to generate NFAT-deficient CD90.1 OTI cells. C57BL/6J mice were purchased from Janvier (Le Genest Saint Isle, France). All mice were housed under SPF conditions and handled in accordance with good animal practices. Typically, 8-12 week old animals were used for infection, bone marrow chimera (BMC) generation and recipients for adoptive T-cells transfer. The responsible state office (Lower Saxony State Office of Consumer Protection and Food Safety) approved all animal experiments performed in Helmholtz-Zentrum für Infektionsforschung, Braunschweig under permit no. 33.19-42502-04-16/2342 and 33.19-42502-04-18/2897.

### Cell lines and Viruses

M2-10B4 (CRL-1972) cells were purchased from American Type Culture Collection (ATCC). C57BL/6 primary mouse embryonic fibroblast (MEF) cells were prepared in-house from C57BL/6J mice. M2-10B4 and MEFs were maintained in DMEM supplemented with 10% fetal calf serum (FCS), 2 mM L-glutamine, 100 IU/mL penicillin and 100 μg/mL streptomycin.

MCMV is derived from pSM3fr-MCK-2fl clone 3.3 BAC (36). MCMV-ie2-SIINFEKL was generated by ‘en passant’ mutagenesis as described previously (37) from the BAC-derived mouse cytomegalovirus pSM3fr-MCK-2fl clone 3.3. SIINFEKL peptide was inserted at C-terminus of *ie2* ORF in MCMV genome. The viruses were reconstituted by BAC transfection on MEF cells. After reconstitution, the virus was propagated on M2-10B4 cells. Virus stocks were prepared according to previously described protocol (38). In brief, virus was pelleted from supernatants of infected cells (26000 x g for 3.5 h). Subsequently, the pellet was re-suspended in VSB buffer (0.05 M Tris-HCl, 0.012 M KCl, and 0.005 M EDTA, adjusted to pH 7.8) and then purified by centrifugation through a 15% sucrose cushion in VSB buffer (53000 x g), and a subsequent slow centrifugation step (3000xg, 5 min) to remove cellular debris. Animals were infected by injecting 10^6^ plaque forming units (PFU) of MCMV via intraperitoneal route.

### *In vitro* virus titration

Animal organs were isolated and homogenized in DMEM medium supplemented with 5% FCS. MEFs were infected with organ homogenates diluted in DMEM. The infection was enhanced by centrifugation for 30 minutes at 1000 x g. The plates were incubated for another 30 minutes at 37°C, 5% CO2. The organ homogenates were then removed, and cells layered with DMEM supplemented with 5% FCS and methylcellulose. The plates were incubated for 4 days at 37°C, 5% CO2 and the plaques were quantified by visual inspection under inverted microscope.

### Bone marrow chimera generation

B6 CD45.1 mice were crossed with C57BL/6J to generate F1 CD45.1^+/-^CD45.2^+/-^ animals. Bone marrow cells were harvested from *Nfatc1^fl/fl^.Cd4cre*, *NFATc2^-/-^*, *Nfatc1^fl/fl^.Cd4cre.Nfatc2^−/−^* (NFAT DKO), WT littermates and CD45.1^+/-^CD45.2^+/-^ animals by flushing femur and tibia bones with FACS buffer (PBS containing 2.5% FCS and 2mM EDTA) and passed through a 70-μm cell strainer. ACK lysis buffer (168 mM NH_4_Cl, 10 mM KHCO_3_, 0.1 mM EDTA) was used to lyse erythrocytes from bone marrow cell suspension followed by T cells depletion using mouse CD3ε MicroBead Kit (Miltenyi Biotec) according to manufacturer’s instructions. Different NFAT-deficient or WT bone marrow cells were mixed 1:1 with CD45.1^+/-^CD45.2^+/-^ WT control bone marrow cells. A small aliquot of the mixed suspension was analyzed with flow cytometry to ensure 1:1 mixing of bone marrow cells. C57BL/6J host animals were exposed to lethal total body irradiation (850 cGray) for myeloablation followed by intravenous injection of 2-4−10^6^ mixed bone marrow cells. BMC animals were infected with MCMV 3 months after reconstitution.

### Flow Cytometry

Spleens and lymph nodes were collected and mashed through a 70-µm cell strainer to generate a single-cell suspension, followed by erythrocytes lysis with ACK buffer. Peripheral blood samples were collected by retro-orbital bleeding and were lysed directly. Before lungs isolation, lungs were perfused via the right heart ventricle with approximately 5 mL PBS to remove circulating blood. The lungs were cut into small pieces and digested for 30 min at 37°C with 0.9 mg/mL collagenase type IV, 0.2mg/mL dispase and 40 ug/mL DNase I in complete RPMI medium (RPMI-1640 supplemented with 2.5% FCS, 100 U/ml penicillin and 100 U/ml streptomycin). Finally, the mononuclear cells were purified by gradient centrifugation over 30% Percoll.

Cells were incubated with anti-CD16/CD32 for 30 min and then washed with FACS buffer. Afterwards, cells were stained for 30 min at 4 °C in the dark with a panel of fluorophore-conjugated antibodies assembled from the following list: anti-CD3 (clone 17A2), anti-CD4 (clone GK1.5), anti-CD8 (clone 53- 6.7), anti-CD11a (clone 2D7) anti-CD27 (clone), anti-CD44 (clone IM7), anti-CD45.1 (clone A20), anti-CD45.2 (clone 104), anti-CD62L (clone MEL-14), anti-CD90.1 (clone OX-7), anti-CD90.2 (clone 53-2.1), anti-CD127 (Clone A7R34), anti-CXCR3 (Clone CXCR3-173), anti-CX3CR1 (Clone SA011F11), anti-KLRG-1 (clone 2F1) and anti-TCRb (clone H57-597). After washing with FACS buffer, the cells were resuspended in FACS buffer containing 7AAD and analyzed using LSR-Fortessa (BD Biosciences) flow cytometer. FlowJo (v.9.6 and v.10.4; Becton Dickinson) were used for data acquisition and analysis.

### Peptide stimulation assay

The SIINFEKL peptide (H-2K^b^-restricted) was synthesized and HPLC purified (95% purity) at the HZI peptide-synthesis platform. For intracellular staining, single cell suspensions were incubated with 1 µg/ml SIINFEKL peptides in complete RPMI medium for 1 h at 37°C followed by brefeldin A (Golgiplug; BD Pharmingen) addition and further 5 h incubation. For intracellular staining, surface-labelled cell suspensions were fixed using eBioscience Foxp3/Transcription Factor Staining Buffer Set or eBioscience IC Fixation Buffer (both from Thermo Fisher). Cell suspensions were stained with following antibodies at 4°C for 30 min: anti-GzmB (clone QA16A02), anti-IFN-γ (clone XMG1.2) and anti-TNF-α (clone MP6- XT22). After washing with FACS buffer, the cells were analyzed using flow cytometer.

### Multimer/Tetramer Staining

Following fluorophore conjugated pMHC tetramers were kindly provided by R. Arens: M45-specific (H- 2D^b^ restricted peptide HGIRNASFI), M57-specific (H-2K^b^ restricted peptide SCLEFWQRV), M38-specific (H-2K^b^ restricted peptide SSPPMFRV) and m139-spcific (H-2K^b^ restricted peptide TVYGFCLL). After incubation with anti-CD16/32, single cell suspensions were incubated with fluorophore conjugated tetramers for at least 30 min at room temperature. Subsequently, the cells were surface stained with fluorophore-conjugated antibodies and analyzed with flow cytometry.

### Adoptive T cells transfer

OT-I cells were isolated from spleen of CD90.1 OT-I mice using Naive CD8a^+^ T Cell Isolation Kit (Miltenyi Biotec) according to manufacturers’ guidelines. 10^4^ naïve NFATc1 KO, NFATc2 KO, NFATc1c2 DKO or WT OT-I cells were adoptively transferred into 8-12 week old recipient animals via tail vein injection. Next day, the animals were infected intraperitoneally with 10^6^ PFU of MCMV-ie2-SIINFEKL.

### RNA isolation and transcriptional analysis

OTI cells were sorted from splenocytes and LNs at 7 days post infection (dpi), and total RNA was extracted using RNeasy Plus Mini kit (Qiagen) according to manufacturer’s instructions. Quality and integrity of total RNA was controlled using 5200 Fragment Analyzer System (Agilent Technologies). The RNA sequencing libraries were generated using NEB Next Single Cell/Low Input RNA Library Prep Kit for Illumina (NEB) according to manufacturer’s protocol and sequenced on a NovaSeq 6000 sequencer (Illumina) using NovaSeq 6000 S1 Reagent Kit (100 cycles, paired end run 2 × 50 bp) with an average of 2.5 × 10^7^ reads per sample. The GEO accession number for all RNA-seq data reported in this paper is GSE228527. Read quality of sequenced libraries was evaluated with FastQC. Sequencing reads were aligned to the reference mouse genome assembly GRCm38 using hisat2 (39). Reads aligned to annotated genes were quantified with featureCounts (40). Read counts were further processed with DESeq2 for quantification of differential gene expression (41). A log-fold change bigger than one and a false discovery rate cut-off of 5% was employed to select significantly over-and under-represented genes. Geneset enrichment analysis was performed using fgsea and clusterProfiler R packages. Analysis was performed on DEGs (p-val < 0.05) from the comparison between NFAT KO and WT OTI cells. The number of permutations was set to 1000.

### Statistical analysis

Comparisons between two groups were performed using the Mann-Whitney U test (two-tailed). Statistical analyses were performed with GraphPad Prism 6-9. p-values < 0.05 were considered significant (*p < 0.05; **p < 0.01; ***p < 0.001), ****p < 0.0001).

## RESULTS

### NFAT KO animals fail to mount persistent CD8^+^ T cell response during chronic infection

Animals with T cells that lack NFATc1 (*Nfatc1^fl/fl^.Cd4cre,* earlier named *Nfat2^fl/fl^.Cd4cre*), NFATc2 (*NFATc2^-/-^*) or both (*Nfatc1^fl/fl^.Cd4cre. Nfatc2^-/-^*; i.e. DKO) were inoculated with MCMV, whereupon blood CD8^+^ T cell response kinetics were longitudinally monitored to define the persistence of virus-specific T cells. We used peptide-MHC tetramers against immunodominant epitopes of MCMV (16) to label the peripheral blood mononuclear cells (PBMCs), and thus measured CD8^+^ T cell responses via flow cytometry (representative gating strategy in Fig. S1). M45-epitope-specific CD8^+^ T cells responded strongly to acute infection, followed by a rapid contraction phase after the first week (Fig. S2A). NFATc1 KO and NFAT DKO animals showed impaired M45-specific responses at 7 dpi. Inflationary M38-specific responses do not contract, but rather accumulate steadily during chronic infection (16, 42). We observed a highly significant reduction in long-term M38-specific CD8^+^ T cell responses of animals lacking NFAT molecules. The total CD8^+^ T cell numbers were reduced in animals lacking NFATc1 in T cells (Fig. S2A, left panel). Nevertheless, NFAT KO animals showed similar levels of MCMV replication during acute infection, suggesting that acute infection control is not impaired in these animals (Fig. S2B). The latent load of virus was also assessed in relevant sites of virus latency such as the spleen and salivary glands (43, 44), and was not altered in NFAT KO animals over the values seen in WT control animals (Fig. S2C). Thus, the lack of CD8^+^ T cell response or persistence in NFAT KO animals was not due to reduced virus replication or latent persistence.

### NFATc1 signaling is required for persistent CD8^+^ T cell responses

In the conventional *Nfatc2*^-/-^ mice not only T cells are NFATc2-deficient. We generated mixed BMC mice and further created a competitive environment for WT versus NFAT-deficient lymphocytes. Bone marrow cells from NFATc1 KO, NFATc2 KO and NFAT DKO or WT littermates were mixed 1:1 with congenic WT bone marrow cells and transferred to C57BL/6J animals whose hematopoietic cells were ablated by lethal irradiation (Fig. 1A). Animals were infected 3 months post bone marrow transplant and CD8^+^ T cell responses were tracked. The NFAT-deficient (CD45.1-CD45.2+) CD8^+^ T cell reconstitution levels were similar to WT (CD45.1+, CD45.2+) controls (Fig. 1B and 1C). NFATc2 KO CD8^+^ T cells showed a modestly increased frequency among total CD8^+^ T cells in line with previous reports (45), whereas the NFATc1 showed an inverse pattern. The frequency of CD45.2^+/+^ T cells remained stable or showed modest reduction during acute infection (Fig. 1C). However, the fraction of NFATc1 and NFAT DKO CD8^+^ T cell populations was significantly reduced during latency for both epitopes tested, where the contraction occurred by 14 days post infection and the populations were stably reduced thereafter.

**Figure 1.**
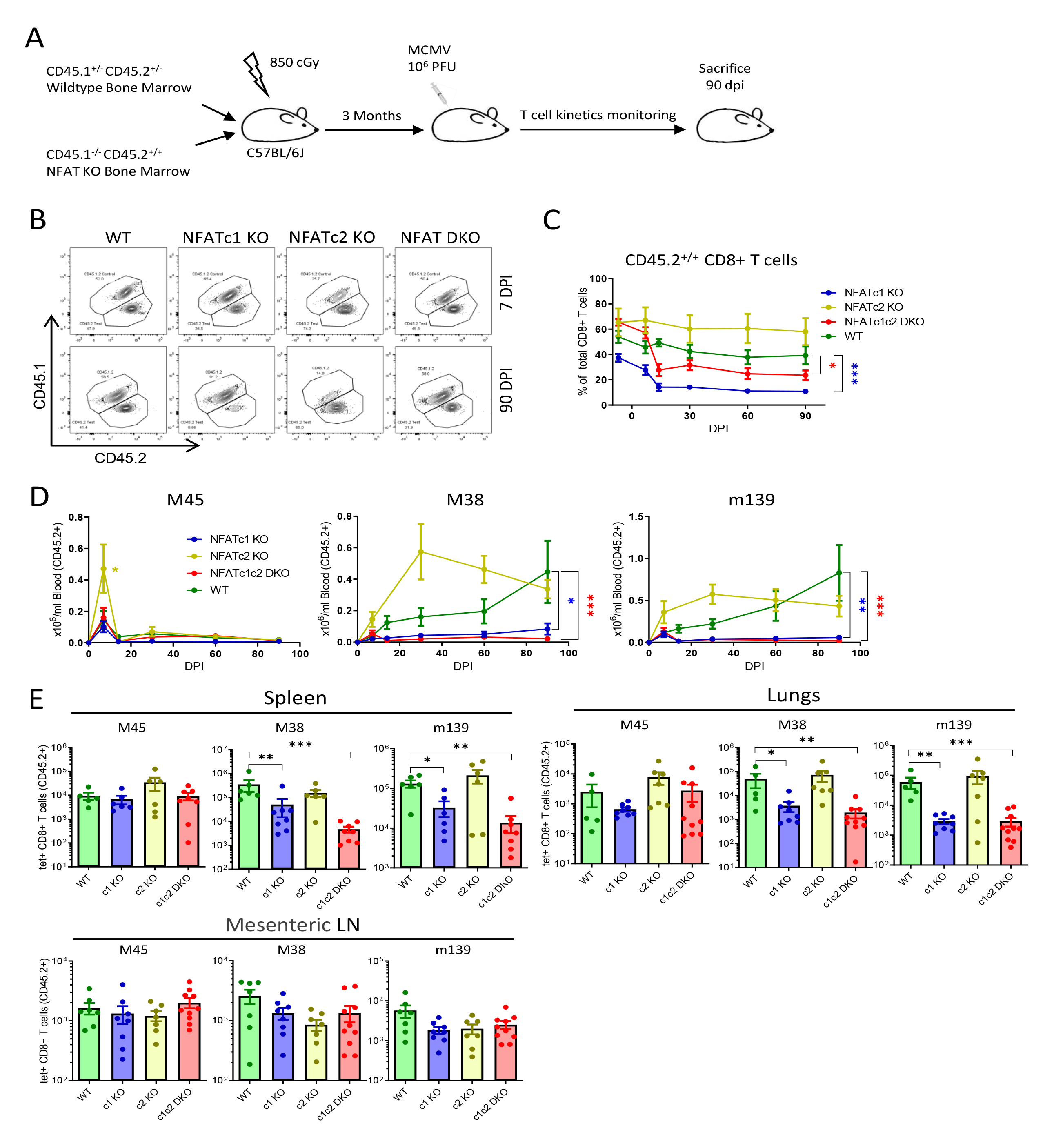
NFATc1 is crucial for persistent CD8+ T cell response during chronic infection. **(A)** Overview of mixed bone marrow chimera (BMC) generation, MCMV infection and CD8+ T cell response monitoring. Lethally irradiated C57BL/6J mice were reconstituted with bone marrow mix (1:1) of the control WT BM (CD45.1^+/-^CD45.2^+/-^) and either NFATc1 KO, NFATc2 KO, NFATc1c2 DKO or WT BM (CD45.1^-/-^CD45.2^+/+^). CD8+ T cell response kinetics were monitored for 90 days following intraperitoneal MCMV infection with 10^6^ PFU. Mice were sacrificed 90 dpi and CD8+ T cell responses in blood and organs were analyzed. **(B)** Representative flow-cytometric plots showing total CD8+ T cells during acute and chronic infection. **(C)** Percentage of the CD45.2^+/+^ subset in the CD8+ T cell population was tracked by labelling peripheral blood. **(D)** CD8+ T cell response was tracked by labelling peripheral blood with MCMV epitope-specific tetramers. Total number of epitope-specific CD8+ T cells among CD45.2+ population is plotted. **(E)** Bar plots represent the mean ± SEM of tetramer+ CD8+ T cells for each epitope in spleen, lungs, and mesenteric LN at 90 dpi. Data are pooled from two experiments and each dot represents one mouse, n=6-10. Statistically significant differences are highlighted; *, p < 0.05; **, p < 0.01; ***, p < 0.001; (Mann-Whitney U Test); mean ± SEM values are plotted.

Interestingly, M45-specific responses in PBMCs of mixed BMC animals were not reduced in absence of NFATc1 (Fig. 1D; Fig. S3A). This was in contrast with the phenotype observed in NFAT KO animals, where a significant reduction in M45-specific cell number was observed in NFATc1 KO and NFAT DKO animals (Fig. S2A). Similarly, NFATc2 KO populations showed a two-fold increase in M45-specific CD8^+^ T cells of BMC mice and no difference in conventional NFATc2 KO animals, which suggests that NFAT signaling in non-CD8 T cells may have contributed to differences in phenotypes in BMC and KO animals. M38 and m139-specific cell responses were significantly reduced in the CD8^+^ T cell populations lacking NFATc1 signaling, because the steady accumulation of antigen-specific cells observed during chronic infection was absent in NFATc1 KO and NFAT DKO cells in absolute (Fig. 1D) and relative terms (Fig. S3A). At times of virus latency, a significant reduction of M38 and m139-specific responses in NFATc1 KO and NFAT DKO cells was observed in spleen and lungs, although similar numbers of antigen-specific T cells were observed across all populations in the mesenteric lymph nodes (LNs) (Fig. 1E). The congenic and co-transferred WT populations in BMC animals (defined as CD45.1^+/-^CD45.2^+/-^) showed consistent levels of CD8^+^ T cell responses during chronic infection regardless of co-transferred NFAT-deficient cells (Fig. S3B). Thus, WT cells mounted normal responses, whereas the T cells that lack NFATc1 signaling were unable to maintain persistent responses in the analyzed tissue compartments. These data suggest that NFATc1 is necessary for maintaining persistent CD8^+^ T cell responses during chronic infection.

### Intrinsic NFAT signaling regulates CD8^+^ T cell differentiation during chronic infection

CD8^+^ T cell persistence is maintained by continuous production of SLECs that are characterized by downregulation of co-stimulatory receptors (CD27 and CD28) and upregulation of killer cell lectin-like receptor subfamily G member 1 (KLRG1) (15, 46). To investigate the phenotype of NFAT KO CD8^+^ T cells during the chronic phase, we used the mixed BMC model. The chronically infected animals were sacrificed at 90 dpi to determine the expression of activation and memory markers on CD8^+^ T cells. NFAT DKO CD8^+^ T cell populations showed reduced expression of CD44 and CD11a and enhanced expression of CD27 and CD62L (Fig. S4). However, among the activated cells (CD44+CD11a+), the majority of NFAT DKO cells presented a memory (Mem) phenotype (KLRG1-CD27+), rather than a SLEC (KLRG1+CD27-) one (Fig. 2A). WT, NFATc1 and NFATc2 KO populations showed a high frequency of CD8^+^ T cells with a SLEC phenotype in PBMCs, spleen and lungs (Fig. 2B). NFAT DKO CD8^+^ T cells, on the other hand, showed an increased frequency of Mem cells. Although the frequency of SLECs in NFATc1 KO population appeared similar to WT, the absolute number of SLEC decreased significantly among NFATc1 KO population in spleen and lungs (Fig. S5A). On the other hand, the majority of the NFAT DKO cells showed a central memory (CM) phenotype (CD62L+CD27+)(Fig. 2C, S5B). The frequency of the splenic CM population was also somewhat increased in NFATc1 KO cells, but this was not statistically significant (Fig. 2C, 2D). Next, we checked the CM phenotype among the virus-specific CD8^+^ T cells. A high percentage of M45-specific cells showed a CM phenotype at 90 dpi in all conditions, which is consistent with the conventional and non-inflationary nature of this epitope (15). In line with previous studies (15, 16), M38- specific CD8^+^ T cells showed predominantly an effector phenotype, but in NFAT DKO mice, the CM frequency was significantly increased (Fig. 2E). The absolute number of CM cells was similar among all groups in different organs (Fig. S5C), arguing that the increase in the frequency of CM in NFAT DKO mice was only relative and due to a reduction of SLEC cells.

**Figure 2.**
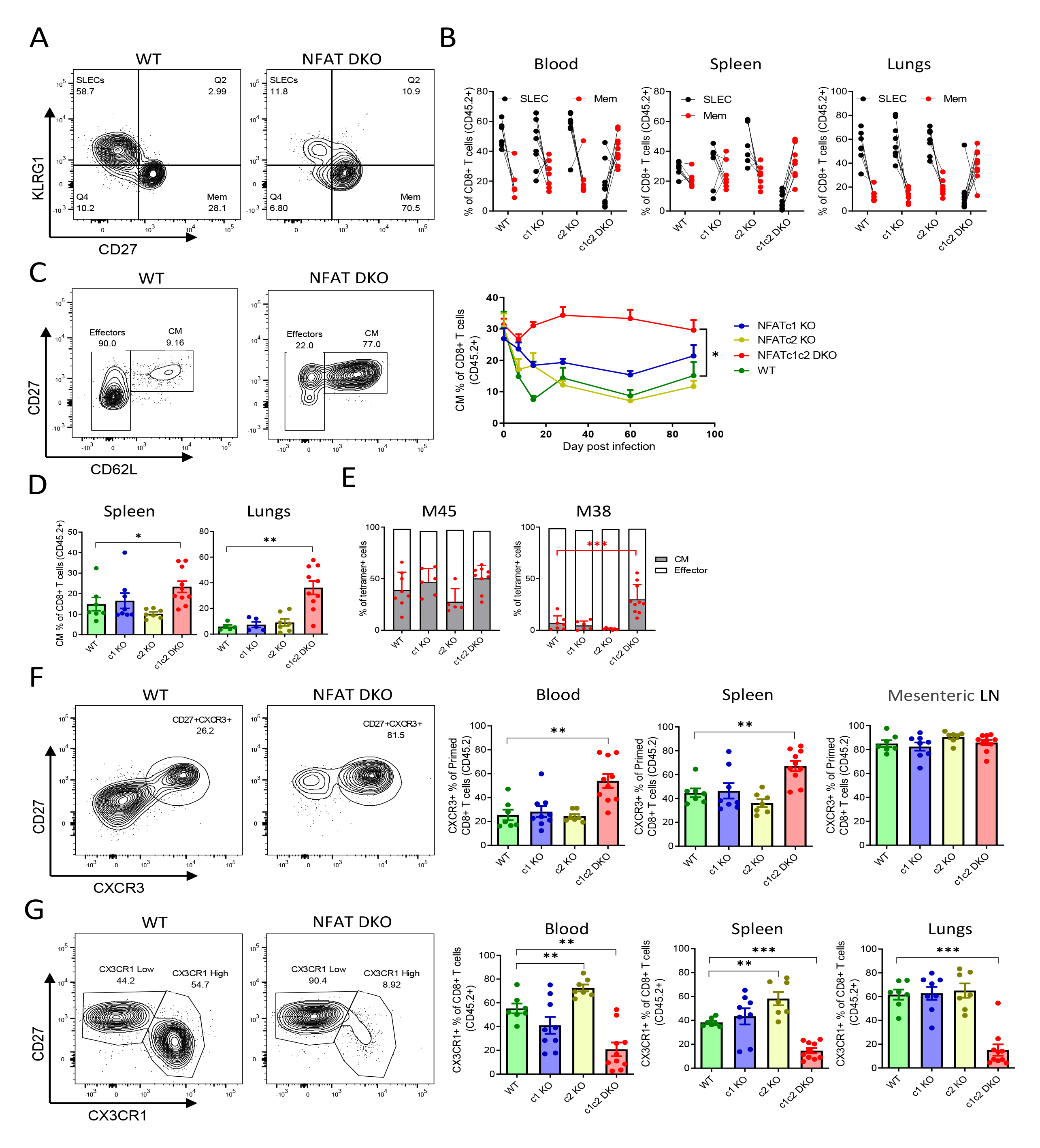
NFAT signaling regulates effector CD8+ T cell differentiation during chronic infection. Mixed BMC animals were infected with 10^6^ PFU of MCMV and analyzed at 90 dpi. **(A)** Representative flow-cytometric plots showing SLECs and Mem populations from BMC mice with WT and NFATc1c2 DKO BM. The plots are pre-gated on primed (CD44+CD11a+) CD45.2+ CD8+ T cells (live CD3+CD8+). **(B)** Pairwise comparison of SLEC and Mem CD8+ T cell frequencies in blood, spleen and lungs of individual mice during chronic MCMV infection. Lines connect data from individual animals **(C)** Flow-cytometric plots showing representative central memory (CM) populations among primed blood CD8+ T cells of chronically infected mice (left). Kinetics of these CM populations in blood are shown on the right Lines connect group means at indicated time points, error bars are SEM. **(D)** Percentage of CM CD8+ T cells in spleen and lungs at 90 dpi. Bat plots represent the group average, error bars are (SEM) each dot is a mouse. **(E)** Percentage of CM cells among M45 and M38 tetramer specific CD8+ T cells from spleen at 90 dpi. Bar plots represent mean ± SEM and each dot is a mouse. **(F)** Representative flow-cytometric plots of blood CD8+ T cells showing CXCR3+ population among primed (CD44+CD11a+) cells (left). Mean CXCR3+ CD8+ T cells population from blood, spleen and mesenteric LN at 90 dpi are shown as mean ± SEM, each dot is a mouse. **(G)** Representative flow-cytometric plots of blood CD8+ T cells showing CX3CR1+ population among primed (CD44+CD11a+) cells (left). Mean CX3CR1+ CD8+ T cells population from blood, spleen and lungs at 90 dpi are shown as mean ± SEM, each dot is a mouse. Data are pooled from two experiments and each dot represents one mouse, n≥7. Statistically significant differences are highlighted; *, p < 0.05; **, p < 0.01; ***, p < 0.001; (Mann-Whitney U Test); mean ± SEM values are plotted.

Since fewer SLECs are generated in absence of CXCR3 on CD8^+^ T cells (47, 48), we next investigated if impaired SLEC generation from NFAT KO populations matched an impaired CXCR3 expression. Interestingly, we observed that NFAT DKO CD8^+^ T cells expressed high levels of CXCR3 in all tested organs (Fig. 2F and Fig. S4), which indicated that NFAT deficiency does not interfere with CXCR3 upregulation upon CD8^+^ T cell activation, but rather with effects subsequent to CXCR3 expression in early T-cell activation. Thus, we analyzed the fractalkine receptor CX3CR1, another important chemokine receptor that has been used as a marker for SLECs. CX3CR1 is upregulated on inflationary CD8^+^ T cells during chronic infections (17, 49), and we observed that NFAT DKO cells did not upregulate CX3CR1 expression (Fig. 2G and Fig. S4), which is in line with the memory-like phenotype observed in the same cells (Fig. 2A, 2B). Similar to the SLEC phenotype, the frequency of CX3CR1+ T cells among NFATc1 KO population was not significantly changed, but the absolute count of the CX3CR1+ CD8^+^ T cells significantly decreased in NFATc1 KO in spleen and lungs (Fig. S5D). These data indicate that the lack of NFAT signaling altered the CD8^+^ T cells differentiation during chronic phase, where the loss of NFATc1 alone led to a significant decrease of SLEC cells, but the loss of both c1 and c2 increased the fraction of cells with a memory phenotype. Thus, the lack of inflationary CD8^+^ T cells responses was associated with altered T cell differentiation.

### Cell-intrinsic NFAT signaling regulates CD8^+^ T cell differentiation following acute infection

Our data suggested that NFATc1 signaling promotes SLEC differentiation and memory inflation during chronic virus infection. In contrast, data from our BMC model showed that the lack of NFAT signaling did not impair early epitope-specific responses in peripheral blood (Fig. 1D). To ascertain in more detail the effect of NFAT signaling in antigen-specific CD8^+^ T cells following acute MCMV infection, we adoptively transferred TCR transgenic OTI T cells on NFATc1 KO, NFATc2 KO, NFAT DKO or WT background to congenic animals that were then infected with MCMV-ie2-SIINFEKL (50). Akin to the BMC model, the lack of NFATc1, NFATc2 or both did not impair CD8^+^ T cell responses (Fig. 3A), but in contrast to it, not only NFATc2 KO, but also NFAT DKO OTI responses were significantly larger than WT response. Absolute numbers of CM OTI cells were increased in absence of NFATc1 or NFATc2, while the relative frequency of CM cells was significantly increased only in absence of NFATc1, but not in absence of NFATc2 (Fig. 3B). The lack of NFAT signaling in OTI cells resulted in reduced SLEC frequency following acute infection (Fig. 3C). The Mem frequency did not change significantly but CD8^+^ T cells with defective NFAT signaling showed a pronounced increase in KLRG1+CD27+ cells, which represent a transitory state between Mem and SLEC. Hence, our data showed that the loss of NFATc1 appears to restrict the transition of the CM phenotype to more mature forms, and this is further enhanced in absence of NFATc2, suggesting non-redundant and distinct functions of NFATc1 and NFATc2.

**Figure 3.**
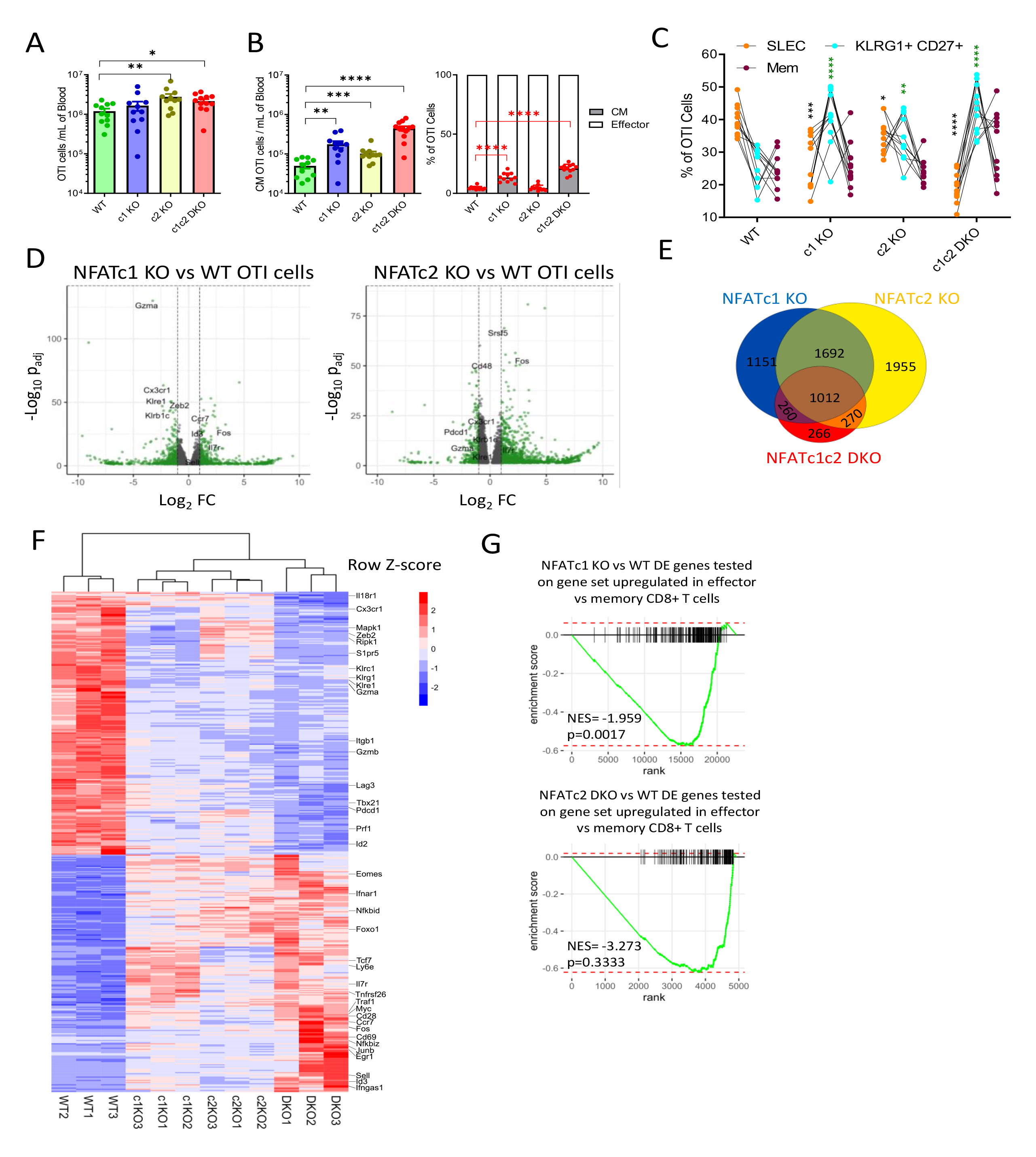
NFAT molecules controls CD8+ T cell differentiation following acute infection and promote distinct transcriptional signature. 10^4^ naïve OTI T cells were transferred to congenic C57BL/6 mice and activated by acute MCMV infection. CD8+ T cell responses were analysed at 7 dpi. **(A)** Absolute number of OTI T cells in blood. **(B)** Absolute and relative size of CM OTI T cell in blood of acutely infected animals. **(C)** Frequency of SLEC, Mem and KLRG1+CD27+ populations among blood OTI T cells are plotted. **(D-G)** Transcriptional analysis (RNA sequencing) was performed on OTI T cells isolated from spleen of acutely infected animals. **(D)** Volcano plot of genes that are differentially regulated in NFATc1 KO and NFATc2 KO OTI cells. **(E)** Venn diagram showing the overlap between differentially expressed genes (>2-fold, <0.1FDR) of different NFAT KO cells compared to WT OTI cells. **(F)** Top 500 genes differentially regulated between NFATc1c2 DKO and WT OTI T cells were selected and their expression in all groups are shown as heatmap. Transcriptional regulators and genes involved in T cell activation have been annotated. **(G)** Negative gene-set enrichment of genes associated with effector CD8+ T cells (62) among differentially expressed genes of NFATc1 KO vs WT OTI T cells and NFATc2 KO vs WT OTI T cells. Panel A-C data was pooled from three experiments, where each dot represents one mouse mean ± SEM values are plotted. Statistically significant differences are highlighted; *, p < 0.05; **, p < 0.01; ***, p < 0.001; ****, p < 0.0001; (Mann-Whitney U Test). The statistical comparisons in panel C are between NFAT KO populations and WT counterpart.

To understand how NFAT signaling regulates CD8^+^ T cell differentiation, we performed detailed transcriptional profiling of NFATc1 KO, NFATc2 KO, NFAT DKO and WT OTI T cells isolated from spleen of acutely infected animals. NFATc1 and NFATc2 KO T cells showed distinct transcriptional signatures with some overlaps (Fig. 3D). We observed a clear under-representation of transcripts related to effector function and SLEC phenotype in NFATc1 KO CD8^+^ T cells. These transcripts code for molecules such as granzyme A (GzmA), Prf1, CX3CR1, KLRG-1 and various other killer-like receptors. Other transcripts that are associated with central memory T cells, such as *Id3*, *Ccr7*, *Il7r* and *Sell*, were significantly upregulated in NFATc1 KO T cells. Transcriptional profile of NFATc1c2 DKO CD8^+^ T cells showed a pattern similar to one from NFATc1 KO T cells (Fig. S6A). Differentially expressed genes (>2-fold, p<0.05) from different NFAT KO cells are shown in a Venn diagram. A total of 4115 and 4929 transcripts changed in NFATc1 and NFATc2 KO T cells, respectively. Among these transcripts, 2704 were affected in both NFATc1 and NFATc2 KO T cells (Fig. 3E). NFAT DKO cells showed only 266 unique transcripts that were not affected in the single KO CD8^+^ T cells. In the heatmap, unsupervised clustering of samples, using top 500 that were differentially expressed (p<0.05) in NFAT DKO, resulted in distinct clades for each genotype (Fig. 3F). NFAT KO cells showed reduced abundance of *Gzma*, *Cx3cr1*, *Klrg1*, and *Prf1*. Some transcripts such as *E2f2*, *Zeb2*, *Id2* were only reduced in NFATc1 and NFAT DKO cells (Fig. 3F and S6B). In contrast, genes involved in memory phenotype like *Id3*, *Eomes*, *Il7r*, *Cd27* and *Ccr7* were selectively upregulated in NFATc1 and NFAT DKO cells. Gene set enrichment analyses confirmed that expression of effector genes (51) was curtailed only in NFATc1 KO, but not in NFATc2 KO cells in a statistically significant manner (Fig. 3G). Taken together, the data showed that NFAT signaling drives transcriptional changes associated with CD8^+^ T cell differentiation and effector function, and NFATc1 and NFATc2 molecules regulate an only partly overlapping set of genes.

### Intrinsic NFAT signaling dictates CD8^+^ T cells trafficking to lymphoid and non-lymphoid organs in early infection

NFAT signaling controlled the CD8^+^ T cell transcription and their differentiation into effector T cells. The expression of chemokine receptors was also affected in CD8^+^ T cells that lack NFAT molecules. This prompted us to investigate if changes in chemokine receptors translate to changes in CD8^+^ T cell migration to different lymphoid and non-lymphoid organs. We investigated the role of NFAT signaling in acute infection of BMC animals, and observed that antigen-specific responses from NFATc1-deficient or NFAT DKO CD8^+^ T cells were impaired in lungs, but not in lymph nodes or spleen (Fig. 4A). Akin to NFATc2 KO OTI responses (Fig. 3A), NFATc2 KO populations in BMC animals showed enhanced CD8^+^ T cells responses to conventional (M45, M57) and inflationary (M38, m139) epitopes in the spleen and blood (Fig. S7A). The NFATc1 and NFAT DKO populations showed reduced antigen-specific responses in lungs, but NFAT DKO CD8^+^ T cells had significantly enhanced responses in mesenteric LN (Fig. 4A). An overall accumulation of NFAT DKO T cells was observed in mesenteric and inguinal LNs (Fig. 4E). The differentiation defects observed during chronic infection in dependence of NFATc1 were also present following acute infection (Fig. 4B-D, S7C).

**Figure 4.**
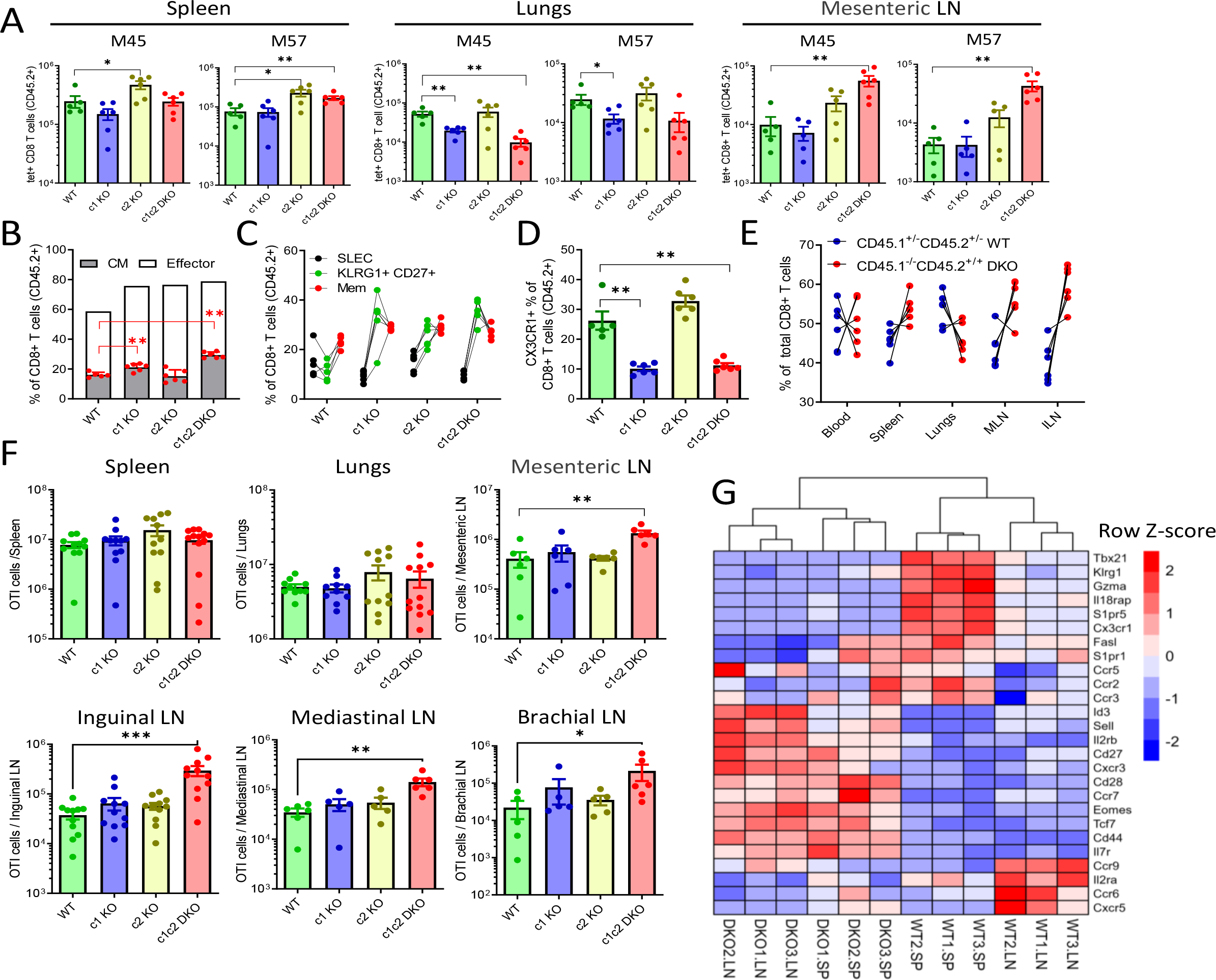
Defective NFAT signaling in CD8+ T cells leads to accumulation of memory CD8+ T cells in LNs. **(A-E)** Mixed BMC animals were infected with 10^6^ PFU of MCMV and dpi CD8+ T cells responses were analysed by tetramer staining and flow cytometry at 7 dpi. **(A)** Total number of indicated tetramer+ CD8+ T cells in spleen, lungs and mesenteric LN. (**B-D**) Relative size of CM, effector **(B)**, SLEC, Mem, KLRG1+CD27+ **(C)** and CX3CR1+ **(D)** CD8+ T cell populations from spleen are plotted. **(E)** Percentages of total CD45.1^-/-^CD45.2^+/+^ NFATc1c2 DKO and CD45.1^+/-^CD45.2^+/-^ WT cells among CD8+ T cells in different organs. **(F-G)** 10^4^ naïve OTI T cells were transferred to congenic animals and activated by MCMV infection. **(F)** Absolute count of total OTI T cells in spleen, lungs and different LNs at 7 dpi. **(G)** WT and NFATc1c2 DKO OTI T cells were isolated from spleen or LNs and transcriptional analysis was performed. Genes involved in T cell activation and migration are shown as a heatmap. Panel A-E show data pooled from two experiments (n=5-6) and panel F have data pooled from 2-3 experiments (n=5-10). Plots show means ± SEM values, each dot represents one mouse. Statistically significant differences are highlighted; *, p < 0.05; **, p < 0.01; ***, p < 0.001; (Mann-Whitney U Test).

To test if the accumulation of antigen-specific CD8 T cells observed in LN in the BMC model is related to CD8^+^ T cell intrinsic signaling, we performed a similar analysis upon acute infection in NFAT deficient OTI T cell. We observed an accumulation of NFAT DKO T cells in mesenteric LN but not in the lungs (Fig. 4F). We analyzed CD8^+^ T cell responses in peripheral and non-draining LNs, such as brachial or inguinal LN, and observed an increased total number of NFAT DKO cells. Furthermore, NFAT DKO cells showed a CM phenotype in all lymphoid and non-lymphoid organs (Fig. S7D). RNA-seq of WT and NFAT DKO OTI cells from LN was compared to the transcriptional profile of cells isolated from the spleen. WT cells isolated from LN and spleen clustered in distinct clades (Fig. 4G), where cells isolated from spleen showed enhanced expression of genes involved in effectors function, such as *Tbx21*, *Klrg1* and *Gzma*. NFAT DKO cells from spleen and LN showed much less differences from each other. In both cases, they failed to upregulate genes involved in effector functions, and showed an increased expression of genes associate with memory phenotype. Furthermore, NFAT DKO cells showed altered expression of genes involved in CD8^+^ T cells migration, such as S1pr1, S1pr5 and chemokine receptors (Fig. S7E). Since memory cells preferably home to lymphoid organs (52, 53), it is likely that the accumulation of NFAT DKO cells in LNs is due to their memory phenotype.

### Virus control by NFAT deficient CD8^+^ T cells

NFATc1 has been shown to control CD8^+^ T cell cytotoxicity (54). Flow cytometric analysis of *in vitro* SIINFEKL re-stimulated splenocytes obtained from MCMV-ie2-SIINFEKL infected mice showed that combined loss of NFATc1 and NFATc2 results in lower IFN-γ production and reduced frequency of IFN- γ^+^TNF-α^+^ T cells (Fig. 5A). We have shown previously that IFN-γ and TNF-α producing CD8^+^ T cells control MCMV virus replication in co-culture settings (38). While CD8^+^ T cells play an important role in virus control, they are not necessary for control of acute virus infection due to redundant immune effectors (55, 56), which may explain the control of acute virus infection in NFAT-deficient animals (Fig. S2B). To test the antiviral capacity of NFAT-deficient CD8^+^ T cells, we used a model where CD8^+^ T cells were required for virus control. Hence, we adoptively transferred WT or NFAT-deficient OTI cells to RAG2gc KO animals, lacking T and B cells and harboring only non-functional NK cells, and infected them with an MCMV expressing the target SIINFEKL epitope. The animals receiving OTI cells did not show any significant weight loss upon infection, whereas the control group without OTI cells showed weight loss and some animals succumbed due to virus infection (Fig. 5B). Interestingly, NFATc1 and NFATc2 deficient OTI cells were able to control the virus infection as efficiently as WT OTI cells (Fig. 5C). On the other hand, NFAT DKO cells controlled the virus with a similar efficacy only in the spleen, but virus titers were significantly elevated in SG, liver and lungs, consistent with the reduced NFAT DKO T-cell migration to non-lymphoid organs (Fig. 4E). Altogether, these data show that NFATc1 and NFATc2-deficient CD8 T cells are cytotoxic and may control virus replication *in vivo,* and only a combined loss of NFATc1 and NFATc2 leads to modest reduction in virus control.

**Figure 5.**
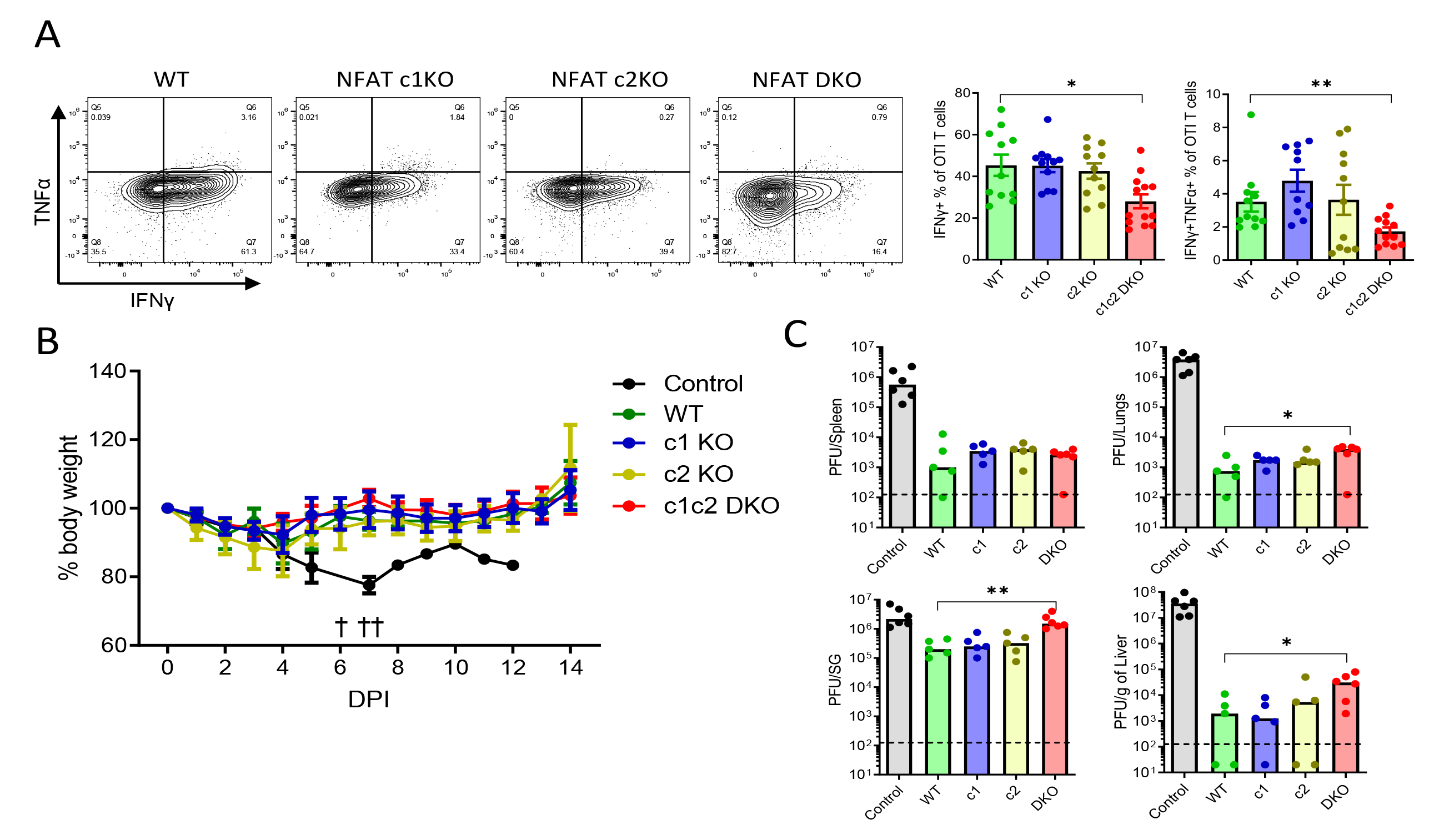
Virus control by CD8+ T cells that lack NFAT signaling. **(A)** 10^4^ naïve OTI T cells were transferred to congenic wildtype animals and activated by MCMV infection. Representative plots show IFNγ and TNFα staining of OTI T cells (FACS plots) and bar plots show IFNγ+ or IFNγ+TNFα+ OTI T cells isolated from spleen at 7 dpi. Error bars show mean ± SEM data pooled from 2 experiments. **(B-C)** 10^5^ naïve OTI T cells were adoptively transferred to RAG2gc KO animals one day prior to MCMV infection with 10^5^ PFU. **(B)** Animal body weight was monitored following MCMV infection and percent of initial body weight is plotted as mean ± SEM. Animals from control group that lost more than 20% of initial body weight were sacrificed, these are denoted by † sign. **(C)** Animals were sacrificed 14 dpi and different organs were collected for virus titration. Virus was titrated from spleen, liver, lungs and salivary glands on mouse embryo fibroblast cells. For control group the values correspond to virus titer at 14 dpi or at the time of death. Bars show median value and each dot represent one mouse. Data pooled from 2 experiments. Statistically significant differences are highlighted; *, p < 0.05; **, p < 0.01; (Mann-Whitney U Test).

## DISCUSSION

It has been proposed that the persistent response of CD8^+^ T cells to MCMV called memory inflation is continuously renewed and maintained by sporadic antigenic stimulation from the latent virus, where latent virus infection and persistent T cell responses constitute a feedback loop. As the latent virus sporadically replicates and stimulates, the T cells limit virus replication and keep MCMV in its latent form. This model predicts that memory inflation crucially depends on TCR activation and downstream signaling.

In our presented study on the role of NFAT signaling in T cell responses to a latently persisting herpesvirus, we observed a crucial role of NFAT family members in the induction of inflationary memory responses, in their maturation and in cell trafficking. NFAT-family members are core regulators of T cell development and maturation (28, 30, 35), activation (26, 45) and functionality (31, 33, 54), with partly redundant and partly distinct roles. We found that NFATc1 is crucial for maintaining persistent CD8^+^ T cell responses during chronic virus infection. The NFATc2 KO T cells, on the other hand, showed overshooting responses suggesting that NFATc1 and NFATc2 serve a non-redundant, and somewhat opposite, function during chronic infection. It was previously documented that heightened levels of NFATc1/αA, an isoform which is transcriptionally upregulated in a positive feedback loop (32), correlates with T cell exhaustion during chronic LCMV infection (33, 57), now thought to be an essential early player in T cell exhaustion (58). Somehow opposing, we report that NFATc1 maintains the functional persistent responses during the MCMV latency. Surprisingly, the number of virus-specific T cells in the LNs was not affected during chronic infection. It has been suggested that antigen presentation on non-hematopoietic cells in LNs drives the persistent memory responses (20). The unchanged CD8^+^ T cell frequencies in the LNs in conjunction with the differentiation defects during the chronic phase would suggest that the lack of NFATc1 leads to insufficient TCR signaling for further differentiation including upregulation of homing receptors like CX3CR1 and / or impaired signal transduction for maintenance in peripheral organs. Contrary to the chronic phase, even the combined loss of NFATc1 and NFATc2 did not impair CD8^+^ T cell responses to acute MCMV infection, suggesting that other signaling cascades induced by strong inflammatory conditions triggered by MCMV infection can compensate the loss of most NFAT signaling. In line, suppression of CD28/B7 or CD27/CD70 signaling cripples primary CD8^+^ T cell responses to MCMV (22), while CD28/B7 and CD27/CD70 signaling is not important, but also not available during chronic MCMV infection (21), possibly explaining the higher dependence on TCR/NFAT signal transduction for the inflated memory. Taken together, these and previous observations advocate that distinct molecular pathways regulate the early and the late CD8^+^ T cell responses to MCMV. NFAT signaling via TCR ligation is crucial for CD8^+^ T cell persistence during the chronic phase, but weaker TCR signaling due to less NFAT can be compensated by the strong costimulation in the context of acute MCMV infection.

Xu et al. demonstrated that NFATc1 and NFATc2 differentially regulate T cell differentiation upon acute LCMV infection (30). In contrast to MCMV acute infection, the loss of NFAT impairs the virus-specific responses during LCMV infection. These contrasting outcomes may be explained by differences in priming requirements for these two viruses, since priming of MCMV-specific cells is facilitated by ‘signal 2’ pathways, rather than ‘signal 3’ cytokine co-stimulation that dominates responses to LCMV (22). As co-stimulation via CD28, CD27 or similar (signal 2) in MCMV priming activates NFAT-independent pathways like NF-κB and MAPK, it appears reasonable to assume that the initial response to MCMV-antigens may proceed in less NFAT-dependent manners than the LCMV responses.

Interestingly, NFATc1 and NFATc2 played distinct roles in persistence of CD8^+^ T cell responses. The stronger differentiation defect observed in absence of NFATc1 than in the ablation of NFATc2, as well as the strongest effects due to combined loss of NFATc1 and NFATc2, suggest a hierarchy of roles that these transcription factors play in regulating differentiation. Namely, our data suggest that NFATc1 appears to be a key regulator of differentiation towards an effector phenotype, while NFATc2 contributes to it, but was not sufficient for cell differentiation in NFATc1 absence. This is in line with a transient, but robust upregulation of NFATc1/αA in effector CD8^+^ T cells (54). Hence, the DKO cells, in which NFATc2 could no longer partly compensate the loss of NFATc1, completely failed to upregulate effector genes and receptors, whereas chemokine receptors and cell adhesion molecules typically present on memory cells were upregulated. Notably, DKO cells with a central memory phenotype that is associated with homing to secondary lymphoid organs accumulated in both draining and non-draining LNs, showing that this is unlikely due to an egress defect, because a pronounced accumulation was not observed only in draining LNs. Furthermore, the transcriptomic profiles of DKO cells isolated from LNs and spleen were remarkably similar, although this was not the case in WT controls, which further argues that DKO cells from spleen preferentially migrated to LNs, instead of lungs, due to their memory phenotype.

Based on *in vitro* experiments, NFATc1 has been suggested as a core modulator of CD8^+^ T cell cytotoxicity (54). However, we observed that selective ablation of NFAT molecules did not impair the ability of CD8^+^ T cells to control the virus in adoptive transfer settings *in vivo*, where only a combined loss of NFATc1 and NFATc2 showed a modest reduction in virus control. This discrepancy may be a reflection of redundant signaling cascades, especially T-cell co-stimulation by other cells, which again could compensate for the loss of NFATc1 in the *in vivo* situation.

Our previous study evidenced that selective ablation of individual NFAT molecules in BM-co-transplanted CD3^+^ T cells ameliorates graft-versus-host disease (GvHD) while maintaining anti-tumor control (34). In this setting, CD8^+^ T cell responses were intact. Here, we show that selective loss of NFAT members in T cells does not impair at all their anti-viral control, which provides an important avenue for CMV disease management. CMV remains a major cause of morbidity and mortality in organ (12) and allogenic stem cell transplantation (allo-HSCT) recipients (11). Organ transplant recipients are often treated with an allo-transplant of CMV-specific T cells for controlling the virus (59). However, broad-spectrum immune suppression including calcineurin inhibition, required to suppress organ rejection or GvHD, also affect the anti-viral function of adoptively transferred T cells. Our study suggests that an advanced therapy with selective NFAT repression may lower the risk of organ rejection while keeping the anti-CMV CD8^+^ T cell compartment functional. Using selective NFAT inhibitors (60) instead of broad-spectrum immune suppression might therefore improve virus control while ensuring survival of transplanted organs. For the setting of allo-HSCT, we found that CRISPR/Cas9-mediated single NFAT member ablation in BM-co-transplanted T cells is sufficient to ameliorate GvHD (61). When we now extrapolate our new findings we predict that single NFAT ablation will not only preserve the anti-tumor effect (Vaeth, 2015), but also CMV reactivation.

## ACKNOWLEDGMENTS

We gratefully acknowledge Anjana Rao for providing the NFAT deficient mice to FBS. We also thank Ayse Barut and Inge Hollatz-Rangosch for excellent technical assistance, and Robert Geffers and Michael Jarek for support with transcriptome analysis. This work was in part funded by the German Research Foundation (DFG) through grants to LCS and FSB (FOR2830, p5) and by the German Centre for Infection Research (DZIF) through a grant to LCS (TTU 07.819).

---

**Figure S1.**
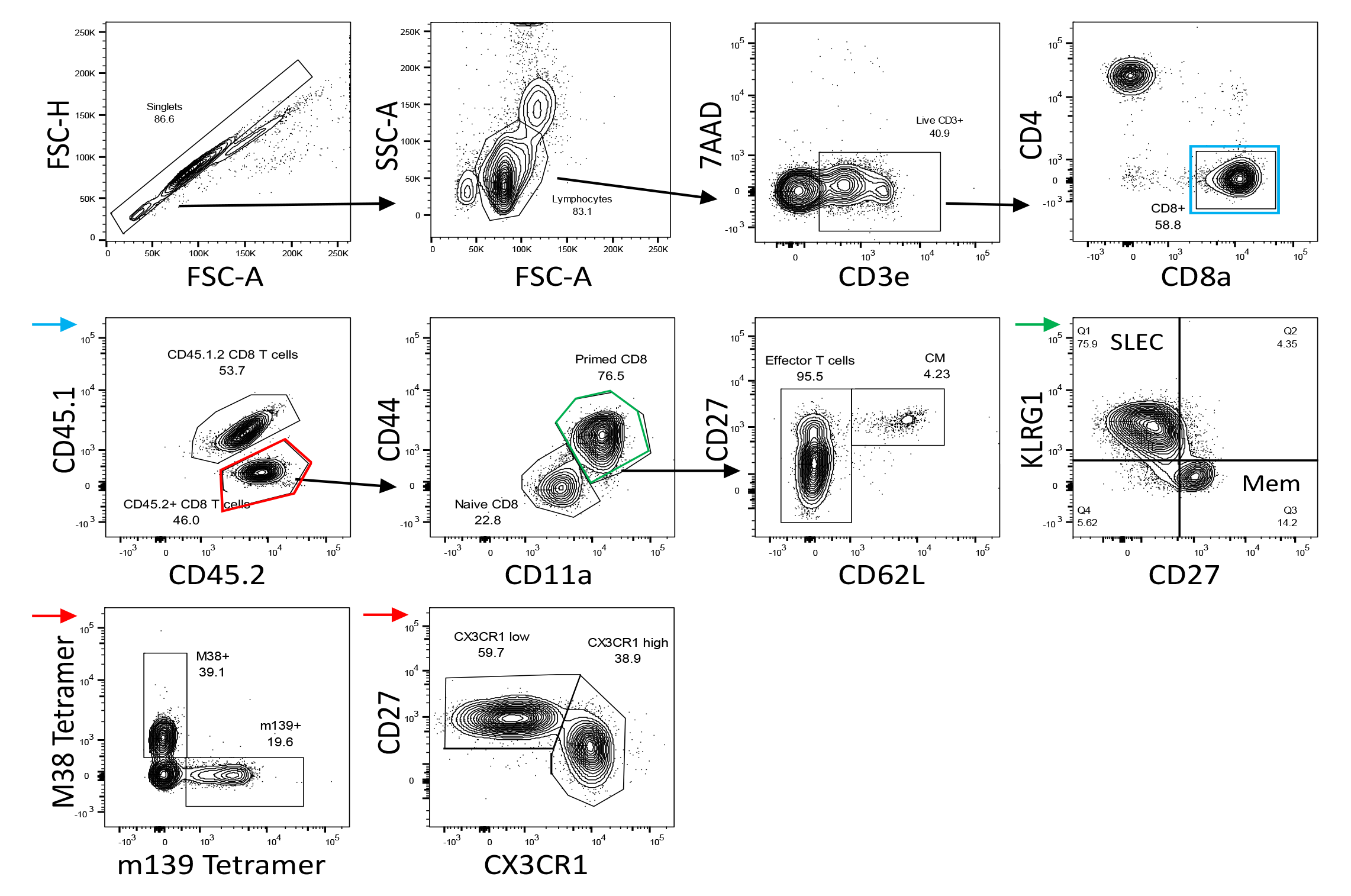
Representative gating strategy. Following singlet gating, lymphocytes were selected with forward/side scatter parameters. Live CD3+ cells were identified by excluding 7AAD stained cells, and CD8+ T cells were selected by CD8a expression. Next, CD8+ T cells (blue gate and arrow) were gated into CD45.2+ single and CD45.1+CD45.2+ double positive populations in mixed bone chimeric animals. Population arising from each bone marrow was progressively gated to define naive and primed CD8+ T cells based on CD44 and CD11a expression. Central memory cells were distinguished from effector cells by CD62L and CD27 expression within primed population. Similarly, short lived effector and memory cell populations (SLEC and Mem) were gated according to KLRG1 and CD27 expression within primed CD8+ T cells (green gate and arrow). Tetramer+ (M38+ and m139+) and CX3CR1+ populations were defined by gating directly on CD45.2+ CD8+ T cells (red gate and arrow) or control CD45.1+CD45.2+ CD8+ T cell populations.

**Figure S2.**
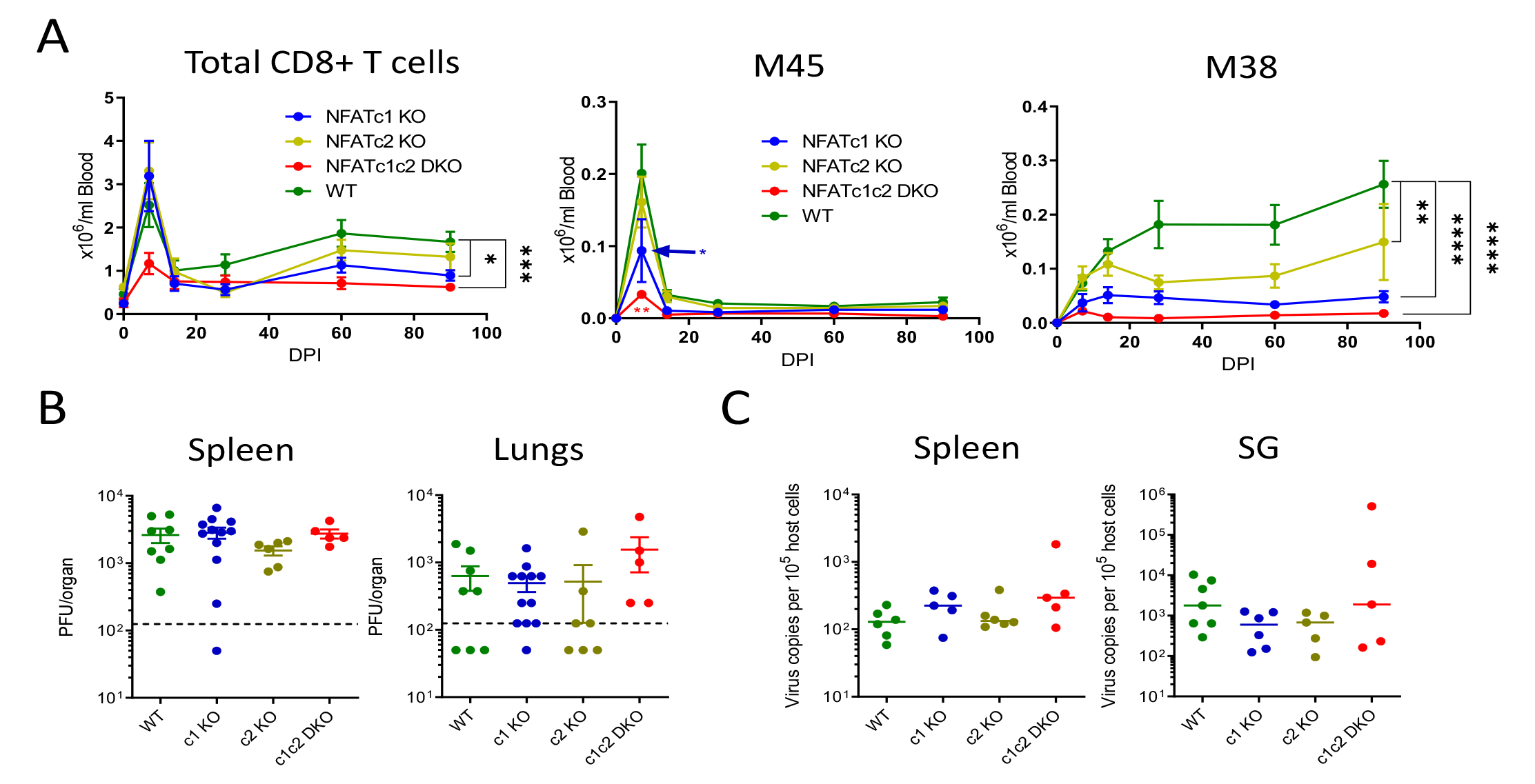
NFAT KO mice fail to mount inflationary CD8+ T cell response. Animals lacking either NFATc1, NFATc2 or both were infected with 10^6^ PFU of MCMV intraperitoneally. **(A)** Total CD8+ T cells and tetramer specific response kinetics in peripheral blood were tracked. Mean ± SEM are plotted for data pooled from 3 independent experiments (n≥12). **(B)** mice were sacrificed at 5 dpi to titrate the virus replication in spleen and lungs. **(C)** Infected mice were sacrificed 6 months post infection to quantify MCMV latent virus load, which is presented as virus copies per 10^5^ host cells in spleen and salivary glands. Data in panel B and C are pooled from two experiments and each dot represent one mouse. Statistically significant differences are highlighted; *, p < 0.05; **, p < 0.01; ***, p < 0.001; ****, p < 0.0001; (Mann-Whitney U Test).

**Figure S3.**
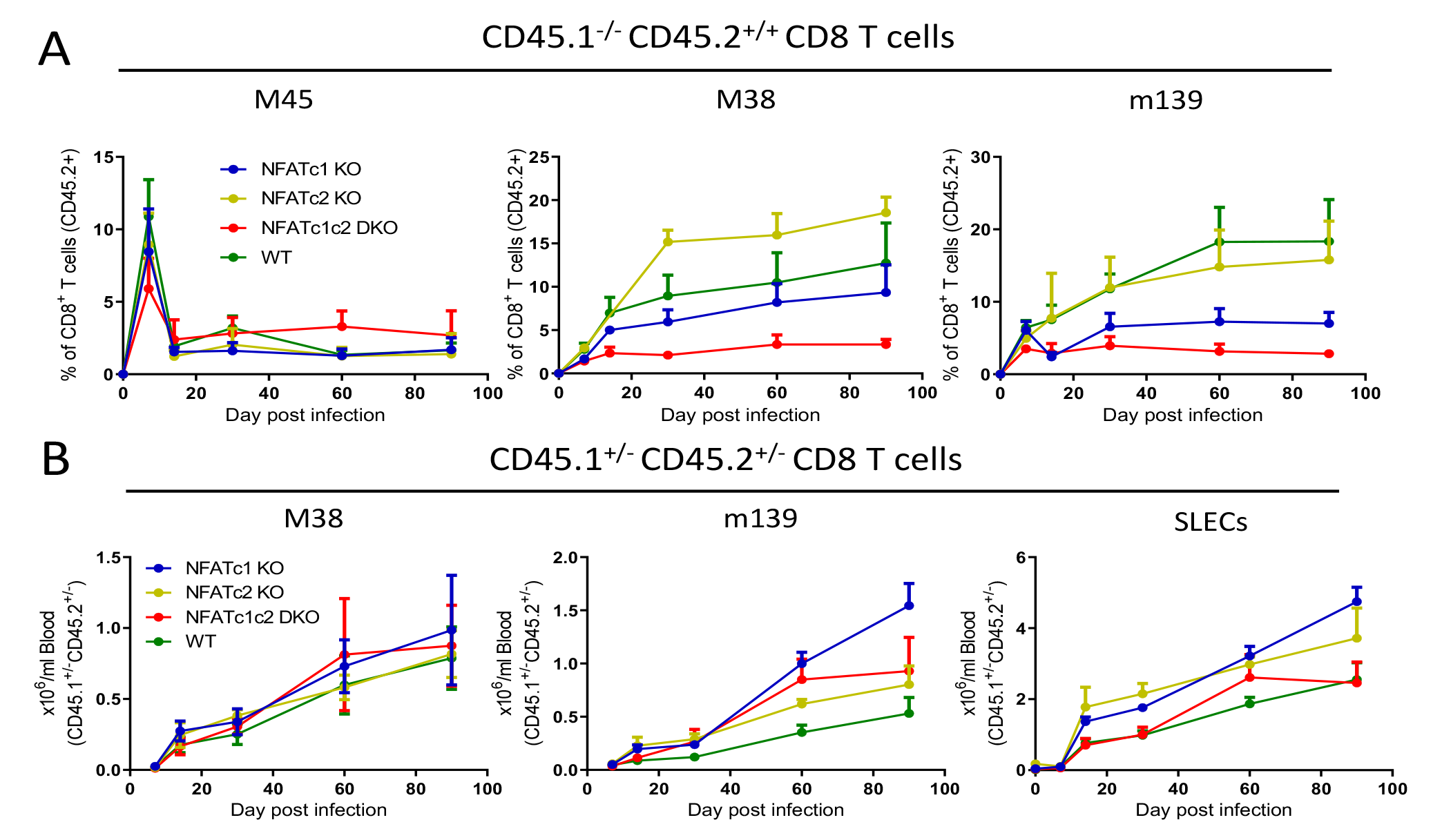
MCMV specific CD8+ T cell responses in mixed bone marrow chimeric mice. Lethally irradiated mice were reconstituted with BM (1:1) of NFATc1 KO, NFATc2 KO, NFATc1c2 DKO or wildtype BM (CD45.2^+/+^) with control wildtype BM (CD45.1^+/-^CD45.2^+/-^). CD8+ T cell response kinetics were monitored for 90 days following intraperitoneal MCMV infection with 10^6^ PFU. **(A)** Relative frequency of tetramer+ cells among CD45.2^+/+^ CD8+ T cells. **(B)** Absolute size of SLEC (KLRG1+ CD27-) and tetramer specific responses from control CD45.1^+/-^CD45.2^+/-^ population. Data are pooled from two experiments and for each group n≥6. Statistically significant differences are highlighted; *, p < 0.05; **, p < 0.01; ***, p < 0.001; (Mann-Whitney U Test); mean ± SEM values are plotted.

**Figure S4.**
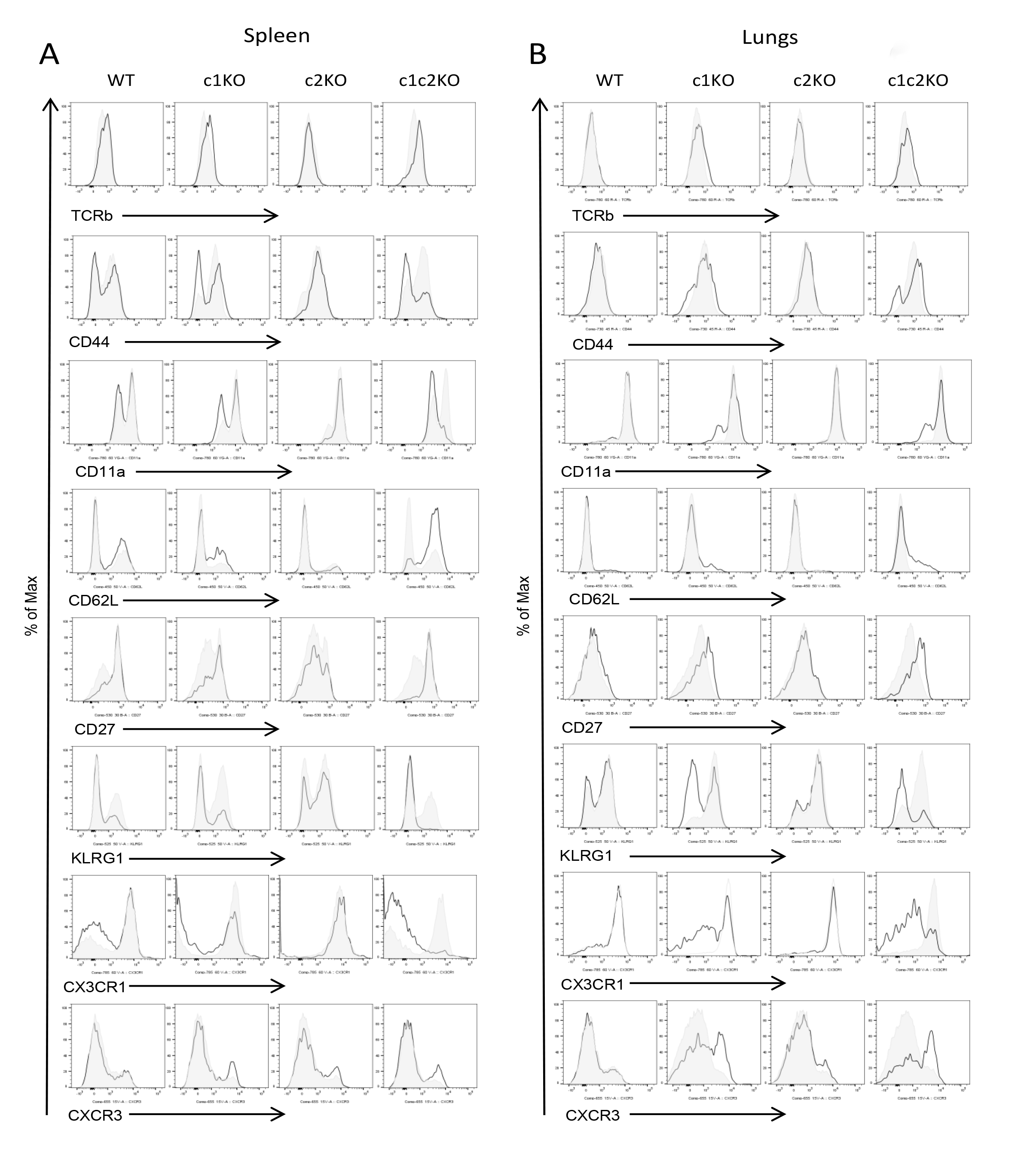
Phenotype of inflationary CD8+ T cells during chronic infection. Lymphocytes from spleen **(A)** or lungs **(B)** from mice infected for 3 months were stained for CD45.1, CD45.2 and CD8 expression as well as the indicated cell-surface molecules. The plots shown are gated on CD45.1^+/-^CD45.2^+/-^ CD8+ T cells (shaded histogram) or CD45.1^-/-^CD45.2^+/+^ CD8+ T cells (black line) from the same sample. Data are representative of at least six individual mice per stain and two independent experiments.

**Figure S5.**
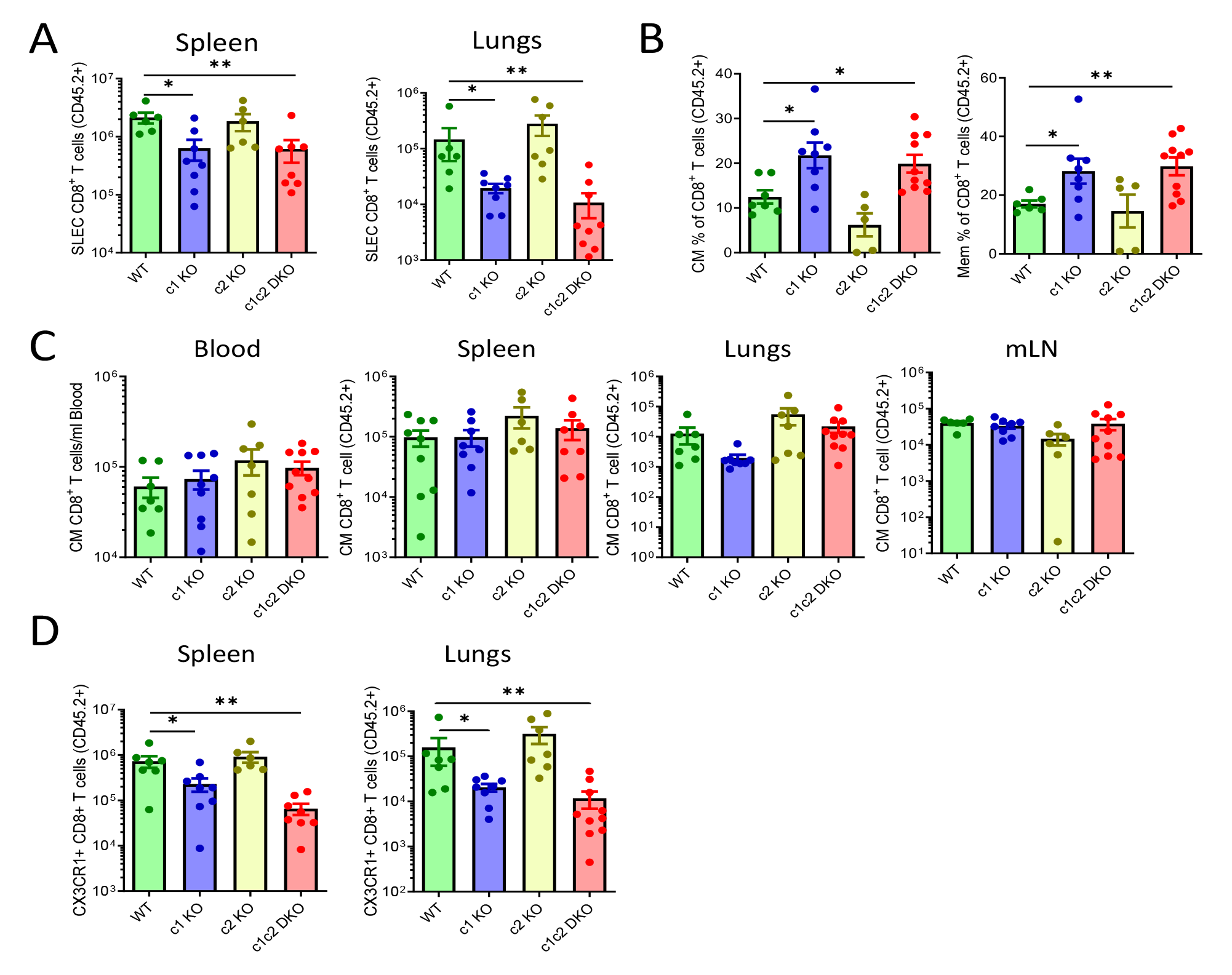
CD8+ T cell differentiation status during chronic infection. Mixed bone marrow chimeric animals were infected with 10^6^ PFU of MCMV and sacrificed at 90 dpi. **(A)** Absolute count of SLECs in spleen and lungs of BMC animals at 90 dpi. **(B)** Frequency of CM and Mem (KLRG1-CD27+) cells among primed CD45.2^+/+^ CD8+ T cells (CD44+CD11a+) from mesenteric LN of chronically infected mice. **(C)** Quantification of CM CD8+ T cells from CD45.2^+/+^ compartment in blood, spleen, lungs and mesenteric LN. **(D)** Absolute count of CX3CR1+ CD8+ T cells in spleen and lungs of BMC animals at 90 dpi. Data are pooled from two independent experiments and each dot represent one mouse. Statistically significant differences are highlighted; *, p < 0.05; **, p < 0.01; ***, p < 0.001; (Mann-Whitney U Test); mean ± SEM values are plotted.

**Figure S6.**
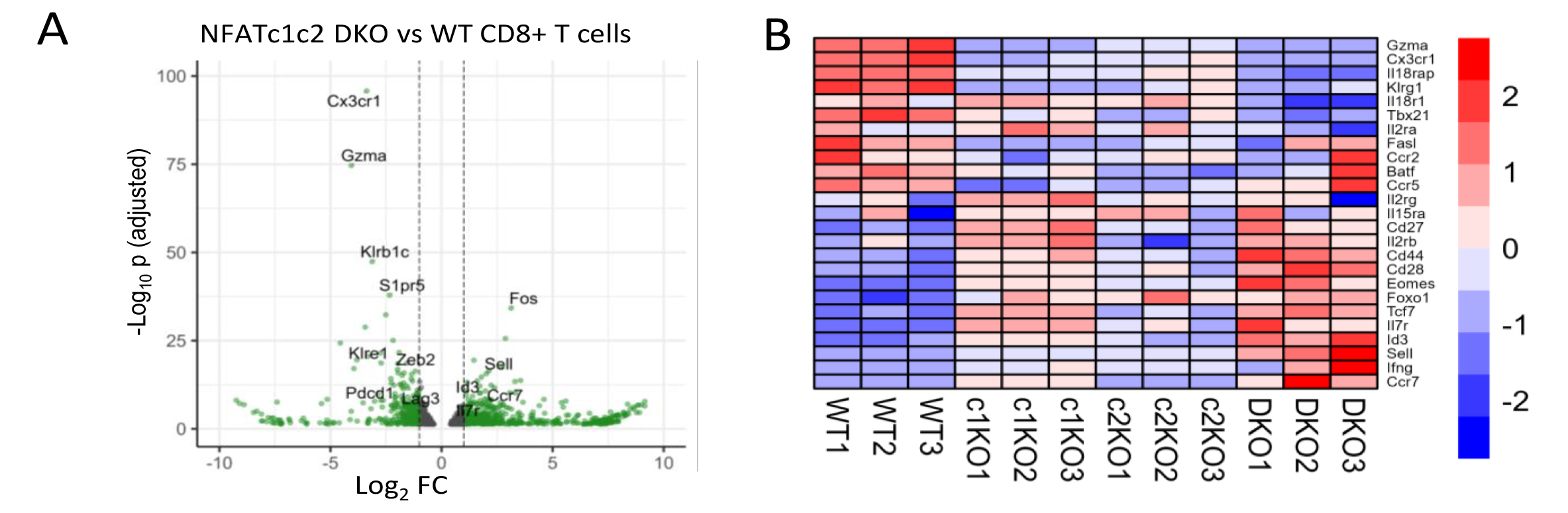
CD8+ T cells lacking NFATc1 and NFATc2 showed transcriptional profile identical to memory CD8+ T cells. Naïve OTI T cells (10^4^) were transferred to congenic animals and activated by acute infection with MCMV. Animals were sacrificed at 7 dpi and CD8+ T cell responses were analyzed. Transcriptional analysis (RNA sequencing) was performed on OTI T cells isolated from spleen of acutely infected animals. **(A)** Volcano plot shows genes that are differentially regulated in NFATc1c2 DKO OTI cells as compared to WT cells. **(B)** Heatmap shows expression of selected genes involved in CD8+ T cell activation and memory differentiation in OTI cells that lack NFATc1, NFATc2 or both. Color shows Z-score differences.

**Figure S7.**
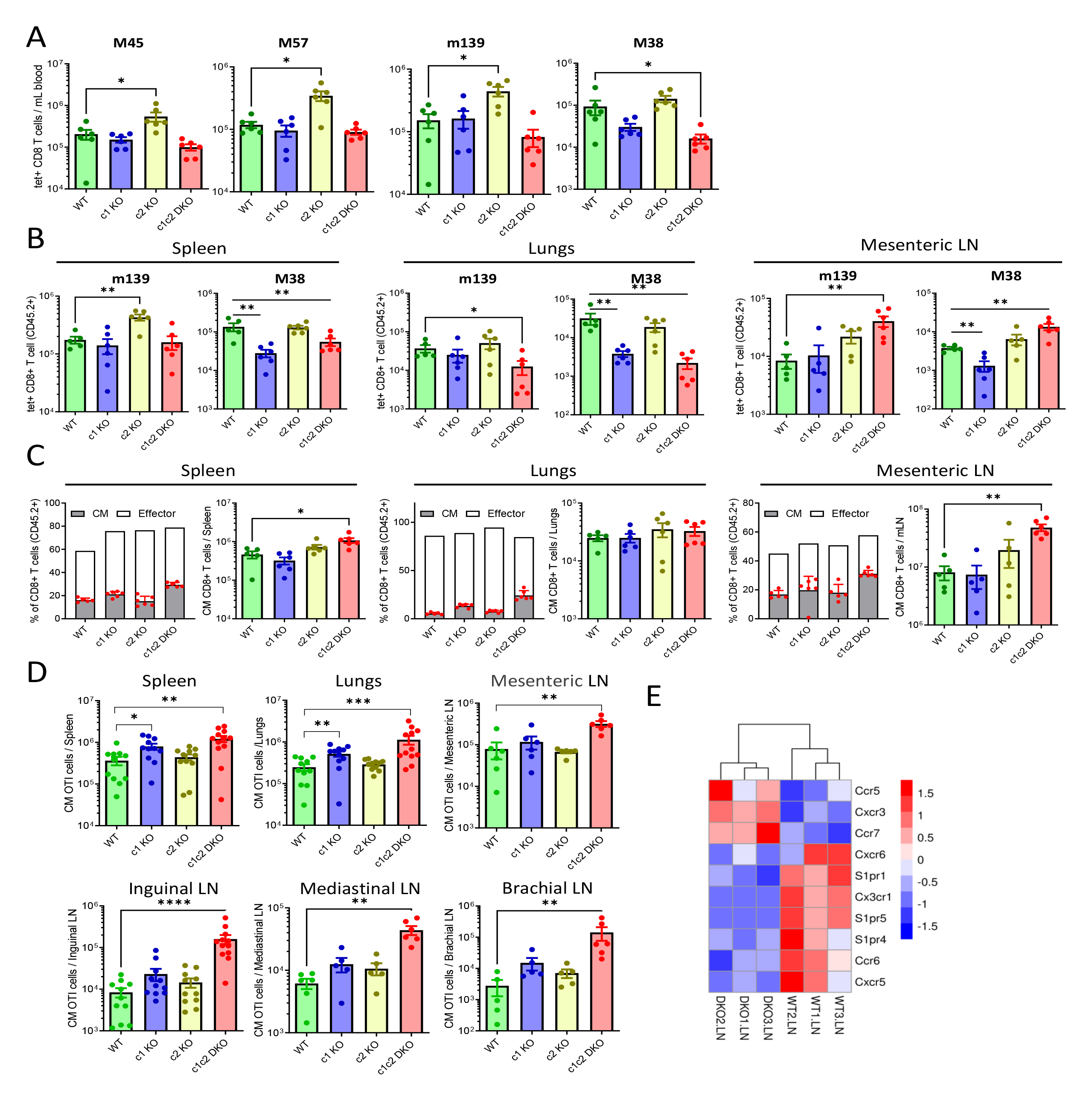
Accumulation of NFATc1c2 DKO CD8+ T cells in Lymph nodes. **(A-C)** Mixed bone marrow chimeric animals were infected with 10^6^ PFU of MCMV and sacrificed at 7 dpi. Absolute count of different tetramer specific CD45.2^+/+^ CD8+ T cells in blood **(A)**, spleen, lungs and mesenteric LN **(B)**. **(C)** Relative and absolute count of CM CD45.2^+/+^ CD8+ T cells in spleen, lungs and mesenteric LN. **(D-F)** 10^4^ naïve OTI T cells were transferred to congenic animals and activated by MCMV infection. **(D)** Absolute count of CM OTI T cells in different organs at 7 dpi is shown. **(E)** Selected genes involved in T cell migration are shown in heatmap. Data are pooled from at least two experiments and each dot represents one mouse; mean ± SEM values are plotted.

